# Evolution of spatial and temporal *cis*-regulatory divergence between marine and freshwater sticklebacks

**DOI:** 10.1101/2022.09.30.510353

**Authors:** Katya L. Mack, Tyler A. Square, Bin Zhao, Craig T. Miller, Hunter B. Fraser

## Abstract

*Cis*-regulatory changes are thought to play a major role in adaptation. Threespine sticklebacks have repeatedly colonized freshwater habitats in the Northern Hemisphere, where they have evolved a suite of phenotypes that distinguish them from marine populations, including changes in physiology, behavior, and morphology. To understand the role of gene regulatory evolution in adaptive divergence, here we investigate *cis*-regulatory changes in gene expression between marine and freshwater ecotypes through allele-specific expression (ASE) in F1 hybrids. Surveying seven ecologically relevant tissues, including three sampled across two developmental stages, we identified *cis*-regulatory divergence affecting a third of genes, nearly half of which were tissue-specific. Next, we compared allele-specific expression in dental tissues at two timepoints to characterize *cis*-regulatory changes during development between marine and freshwater fish. Applying a genome-wide test for selection on *cis*-regulatory changes, we find evidence for lineage-specific selection on several processes, including the Wnt signaling pathway in dental tissues. Finally, we show that genes with ASE, particularly those that are tissue-specific, are enriched in genomic regions associated with marine-freshwater divergence, supporting an important role for *cis*-regulatory differences in adaptive evolution of sticklebacks. Altogether, our results provide insight into the *cis*-regulatory landscape of divergence between stickleback ecotypes and supports a fundamental role for *cis*-regulatory changes in rapid adaptation to new environments.

## Introduction

Understanding how organisms adapt to new environments is a major goal in evolutionary biology. Central to this goal is understanding what genetic changes underlie adaptive traits. Threespine sticklebacks (*Gasterosteus aculeatus*) are a powerful model for studying the genetic basis of adaptation[1]. After the end of the last ice age, marine sticklebacks colonized thousands of freshwater habitats in the Northern Hemisphere [2]. In these freshwater environments, populations have rapidly evolved a number of traits that distinguish them from the ancestral marine form. While adaptation to each lake or stream is independent, several traits have evolved repeatedly across multiple freshwater systems either through parallel, convergent, or distinct genetic changes (e.g., changes in body shape, skeletal armor, dentition, behavior, and pigmentation)[2–6]. The repeated evolution of similar phenotypes in freshwater systems is strong evidence that these traits reflect local adaptation and provide a powerful platform for studying the genetic architecture of adaptive phenotypic evolution [7–9].

Mutations in *cis*-regulatory elements can change how nearby genes are regulated. Such mutations are thought to be an important substrate for adaptive evolution[10–12]. In contrast to protein-coding changes, *cis*-regulatory mutations can alter the expression of gene targets in tissue- or temporally-specific ways. As a consequence, *cis*-regulatory changes may be less constrained by the deleterious side-effects of negative pleiotropy, making this class of mutations important targets for natural selection [10,11]. *Cis*-regulation has been shown to be the major driver of local environmental adaptation in recent human evolution [13], and likewise plays a central role in the local adaptation of sticklebacks to freshwater environments. Genome scans have found that genomic regions associated with recurrent divergence between ecotypes are predominantly intergenic, suggesting parallel divergence may often involve the reuse of pre-existing gene regulatory variation [8]. C*is*-regulatory mutations have been implicated in specific morphological differences between marine and freshwater forms, including the loss of pelvic spines [14], bony armor plates [7], changes in pigmentation[4], and increased pharyngeal tooth number [6,15]. While these lines of evidence suggest an important role for gene regulatory evolution in stickleback adaptation, the global *cis*-regulatory landscape of marine-freshwater divergence remains poorly understood. Exploration of *cis*-regulatory changes between ecotypes has largely been limited to assaying individual gene targets in a small number of tissues (e.g., [6,7,16,17]). Transcriptome-wide *cis*-regulatory divergence between marine and freshwater fish has been characterized in two tissues so far: the gills[18] and ventral pharyngeal tooth plates [19]. Surprisingly, these two tissues showed highly divergent regulatory landscapes[18], suggesting tissue-specific regulatory architecture may play an important role in stickleback adaptation.

Here we survey global *cis*-regulatory divergence between marine and freshwater sticklebacks in seven tissues to understand the role of gene expression evolution in adaptive divergence. To characterize *cis*-regulatory changes between ecotypes, we crossed marine and freshwater fish to generate F1 hybrids. As F1 hybrids carry both a marine and freshwater copy of each chromosome, alleles from both parents are present in the same cellular environment (e.g., subject to the same *trans* factors). Expression differences between the two parental alleles (i.e., allele-specific expression) can therefore only result from *cis*-regulatory changes [20,21]. We use this approach to examine a collection of tissues important for behavioral, physiological, feeding, and morphology differences between marine and freshwater forms (i.e., brain, liver, eyes, flank skin, dorsal and ventral pharyngeal tooth plates, and mandible). As morphological changes include especially dramatic changes to the craniofacial skeleton and dentition[6,22], likely reflecting adaptations to different diets in freshwater, we also examine a second developmental timepoint in dental tissues to characterize *cis*-regulatory modifications during development. We use these data to dissect the landscape of *cis*-regulatory divergence and then ask whether these changes are associated with genomic signals of selection. Overall, our results highlight the importance of the tissue- and developmental stage-specific *cis*-regulatory changes in marine-freshwater divergence and the importance of *cis*-regulatory variation in local adaptation.

## Results and Discussion

### Extensive allele-specific expression across tissues in fresh-water-marine hybrids

To investigate *cis*-regulatory divergence between marine and freshwater individuals, we analyzed allele-specific expression in F1 hybrids between marine and freshwater fish (freshwater Paxton lake benthic [PAXB] x marine Rabbit Slough [RABS])(Figure 1A). Seven tissues were collected from F1 hybrids at the young adult stage (~35 millimeters [mm] standard length [SL]) (brain, eyes, liver, flank skin, ventral pharyngeal tooth plate [VTP], dorsal pharyngeal tooth plate [DTP], mandible) (Figure 1A-B). Additionally, three dental tissues (mandible, VTP, and DTP) were also collected from full-siblings at an earlier juvenile stage (15-20mm SL) for a temporal comparison of dental development (hereafter, “early” vs. “late” developmental timepoint). We sequenced mRNA from each tissue for two biological replicates, obtaining a median of 66.7 million reads per sample (Tables S1, S2). To phase heterozygous sites in F1s, we also performed whole-genome sequencing of the freshwater parent (PAXB) to an average coverage of ~30X (Figure S1).

**Figure 1.**
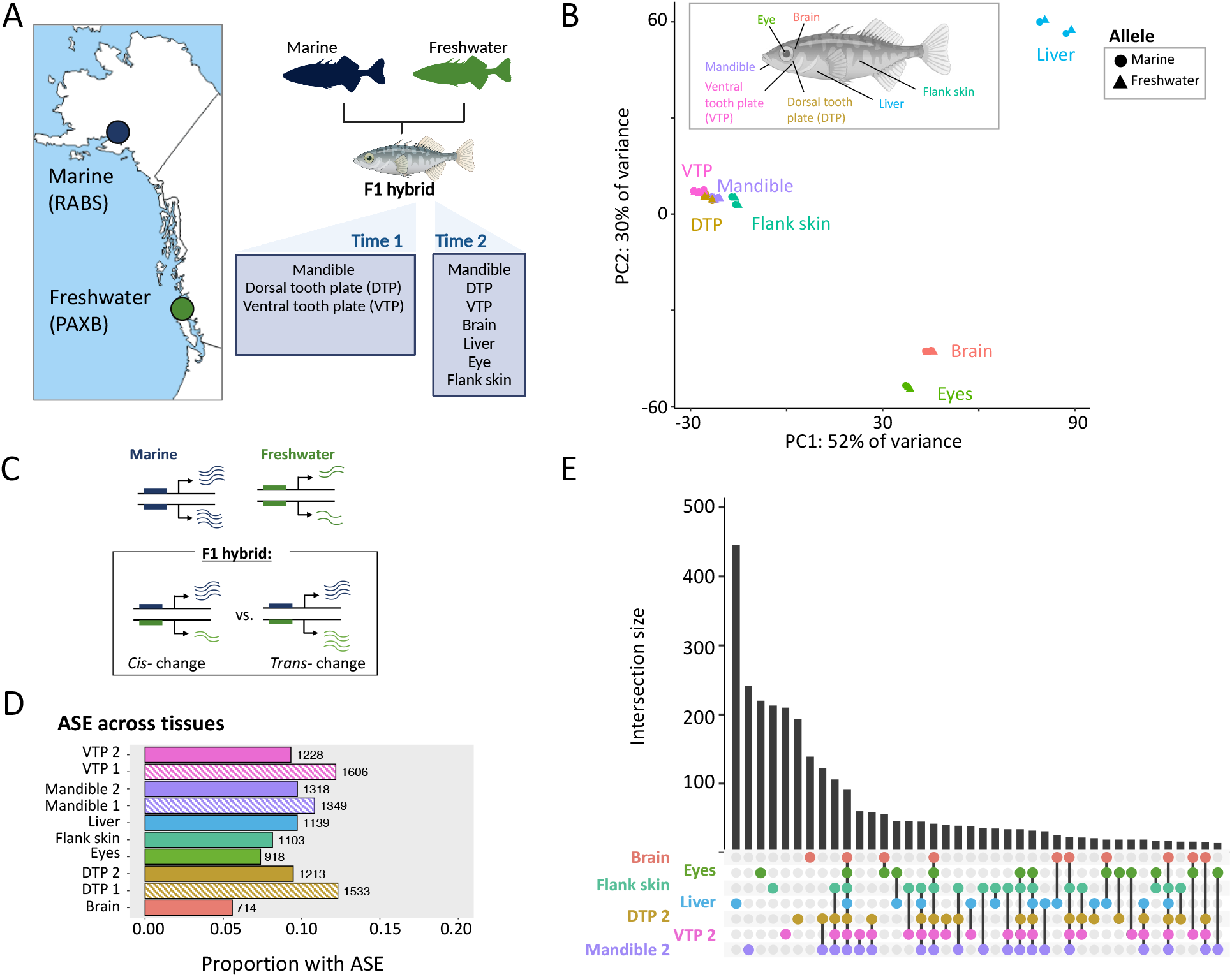
The *cis*-regulatory landscape of divergence between marine and freshwater stickleback. **A**. Marine (RABS) and freshwater (PAXB) sticklebacks were crossed to produce F1 hybrids. Tissues were collected for RNAseq across two developmental timepoints from full siblings. **B**. Principal component analysis of allelic counts from marine-freshwater F1 hybrids. Allelic reads cluster by tissue of origin on PC1 and PC2. **C**. A schematic of how regulatory divergence can be dissected with F1 hybrids. Here, a gene is upregulated in marine fish (wavy lines). In an F1 hybrid, differential expression between the freshwater and marine allele (e.g., allele-specific expression, ASE) indicates a *cis*-regulatory change. In contrast, equal expression of the two alleles indicates a *trans*-only change. **D**. Numbers and proportions of genes with ASE across tissues. Striped bars indicate tissues from the early timepoint. **E**. UpSet plot showing distinct intersections of genes with ASE across tissues from the late timepoint. A single dot in a column indicates ASE specific to one tissue, whereas multiple dots connected by lines indicate ASE shared across multiple tissues.

Principal component (PC) analysis of gene-wise mRNA abundance and allele-specific expression (ASE) revealed tissue-type to be the primary driver of variation (Figures 1B and S2). Allele-specific expression values clustered largely by tissue of origin on PC1 and PC2 (PC1: 52% of the variation, PC2: 30% of variance). Dental tissues formed their own cluster to the exclusion of other tissues, as did eyes and brain. Flank skin, where bony lateral plates develop, also formed a group with dental tissues on PC1 (Figure 1B). PC analysis of allele-specific expression of dental tissue timepoints also separated samples based on developmental stage (early vs. late) on PC1 or PC2 (Figure S3).

Extensive ASE was found across tissues (Figure 1C-D). Nearly 33% of genes (4,411) were found to have significant ASE in at least one tissue or tissue-timepoint (DESeq2 Wald-test, FDR<0.05, see Table S3, Figure 1D; 13,551 genes tested). In each tissue, these ASE genes accounted for approximately 5-12% of genes surveyed. Dental tissues had the greatest number of ASE genes overall, particularly at the earlier developmental timepoint (Figure 1D). The lowest number of ASE genes was identified in the brain (714 genes, 5.6%). The number of ASE genes identified in a tissue was not related to differences in read depth between tissues (Figure S4).

Comparing overlap of genes with ASE between tissues, we found that the largest distinct groups were tissue-specific rather than shared, indicating largely tissue-specific *cis*-regulatory divergence between marine and freshwater fish (Figure 1E). Across the seven tissues sampled at the late timepoint (SL ~35mm), 1,660 genes showed ASE in only one tissue (48% of genes with ASE overall). In particular, the liver was found to have the greatest number of unique ASE genes (366 genes, 32% of genes with ASE in liver). Comparisons between tissues also revealed many genes with evidence for shared ASE (Figure 1E).

In particular, we found high overlap between eyes and brain (29% shared), between dental tissues (36%-44%), and between dental tissues and flank skin (32%-36%)(Table S4). In contrast, few genes (~2%) showed ASE across all tissues (78 genes across all tissues, 92 genes at the late developmental timepoint). For genes with ASE in multiple tissues, directionality was typically maintained, with only 236 genes showing a change in which parental allele was upregulated between tissues.

### Widespread heterogeneity in allele-specific expression across tissues in marine-freshwater hybrids

Comparisons of ASE across tissues revealed abundant *cis*-regulatory divergence between marine and freshwater fish. To investigate variation in allele-specific expression between tissues in marine-freshwater hybrids, we employed a Bayesian approach to partition genes in a tissue into three states – no ASE, moderate ASE, and strong ASE – based on the numbers of reads supporting the marine and freshwater allele [23]. Tissues are further classified as showing ASE heterogeneity if the strength of ASE varied across tissues (e.g., ASE is present in some tissues but absent in others or varies in magnitude between different tissues). Finally, we consider a sub-state of ASE heterogeneity to be tissue-specificity, where the ASE state (i.e., moderate, strong ASE, or no ASE) is observed in only one tissue despite expression of the gene across multiple tissues. Consequently, tissue-specificity describes cases where ASE state is unique to a single tissue.

We found that heterogeneity in ASE between tissues was common. Comparing across the seven different tissues collected at our second timepoint, we found that 44% of genes with ASE were classified as having heterogeneous ASE at a posterior probability (PP)>0.9 (at PP>0.95, 38%)(Figure 2A; Full list in File S1). Nearly all the genes with ASE heterogeneity (99%) did not show ASE in at least one of the tissues surveyed, with the remaining 1% showing evidence for ASE of varying magnitudes across all tissues. Evidence of tissue-specificity was also found for 448 genes (PP>0.9, File S1)(Figure 2B). Liver harbored the greatest number of genes with tissue-specific ASE (141 genes), followed by the eyes (66 genes). Repeating this analysis to incorporate dental tissues from the early timepoint and late timepoint, we also identified 73 genes with developmental- and tissue-specific ASE in tissues from the early developmental stage (File S1). Overall, allele-specific expression across tissues was found to be highly heterogeneous, likely reflecting tissue-specific *cis*-regulatory differences between marine and freshwater individuals.

**Figure 2.**
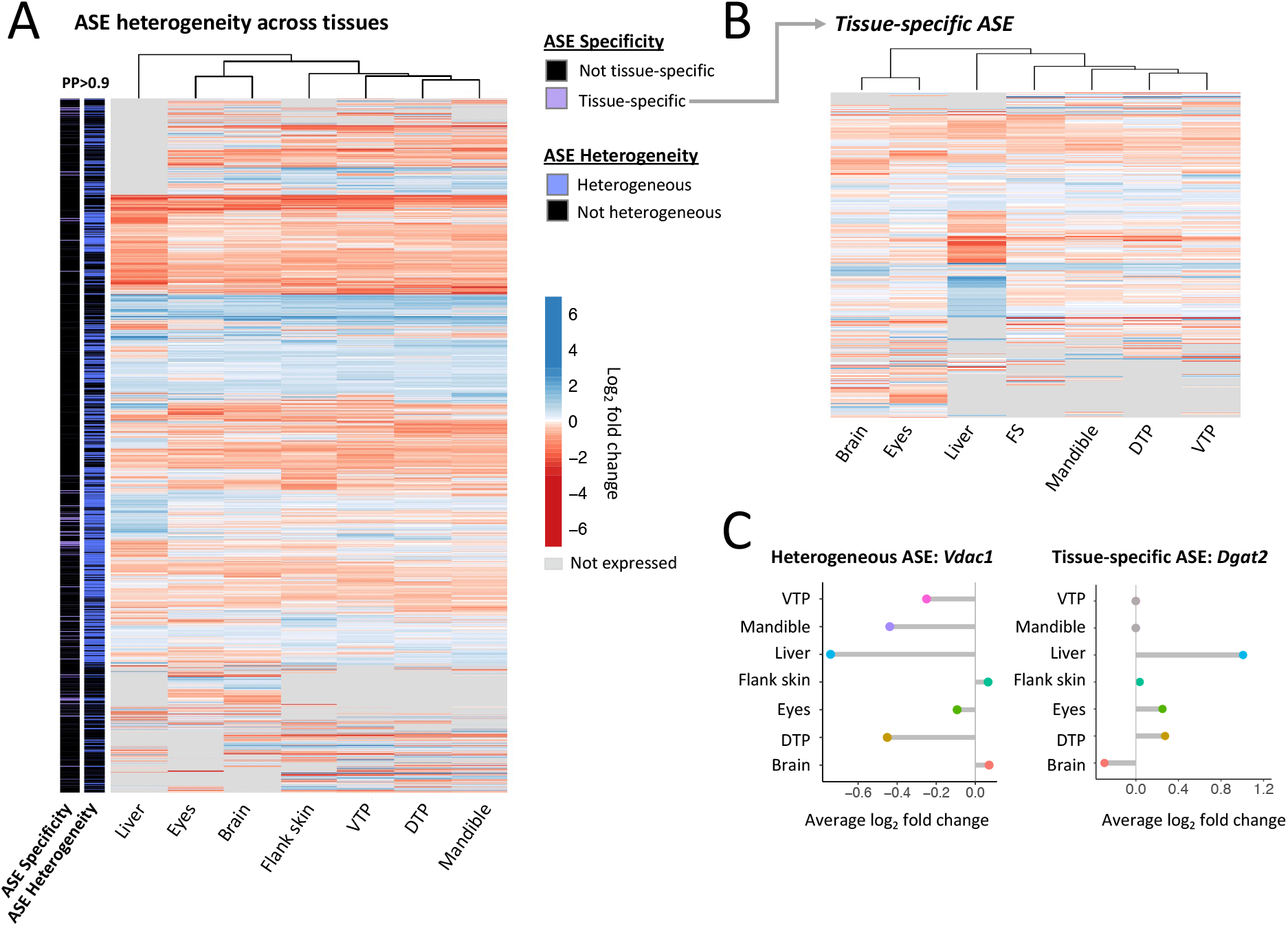
Heterogeneity of allele-specific expression across tissues. **A.** A heatmap of genes with evidence of allele-specific expression in late development. The left bars indicate genes where ASE varies across tissues (in presence or magnitude, “ASE Heterogeneity”) and substate of ASE heterogeneity where ASE patterns are specific to one tissue (tissue-specific ASE, “ASE Specificity”) at a posterior probability of >0.9. Genes are colored by average log2 fold change in each tissue. Gray panels indicate the gene is not expressed in a given tissue. **B**. Heatmap of genes with evidence for tissue-specific ASE. **C**. Examples of genes with heterogeneous ASE (*Vdac1*, left) and tissue-specific ASE (*Dgat2*, right). ASE was observed for voltage-dependent anion channel *Vdac1* in some tissues (e.g., liver, DTP2, mandible) but not others, where triglyceride synthesis gene *Dgat2* only shows evidence for ASE in liver.

Several genes with tissue-specific ASE were of interest for their reported tissue-specific functions in other systems (File S1). For example, *Dgat2* was expressed in five tissues at the second timepoint but found to have liver-specific ASE (Figure 2C). *Dgat2* is involved in triglyceride synthesis and plays an important role in energy metabolism; in mammals and zebrafish, gene mutants are associated with fatty liver [24,25]. In dental tissues, genes with tissue-specific expression include a number of genes involved in tooth and bone formation (e.g., *Spp1, Dlx1a, Odam, Sox2, Epha3, Ssuh2rs1, Tgfbr2b, Stc2a*)(File S1). *Stc2a*, which was found to have tissue-specific ASE in the early mandible, was also recently shown to underlie changes in pelvic spine length between stickleback populations [16].

### Temporal differences in allele-specific expression during dental development

Marine and freshwater sticklebacks show a number of phenotypic differences associated with feeding morphology (e.g., larger jaws, more teeth), likely reflecting adaptations to larger prey found in the benthic zone of lakes [2,26]. Divergence in tooth number arises during late development, providing an opportunity to study *cis*-regulatory divergence in the context of developmental evolution [6,19,27]. To investigate *cis*-regulatory divergence during dental development, we examined ASE in three dental tissues at two developmental timepoints (Figure 3A).

**Figure 3.**
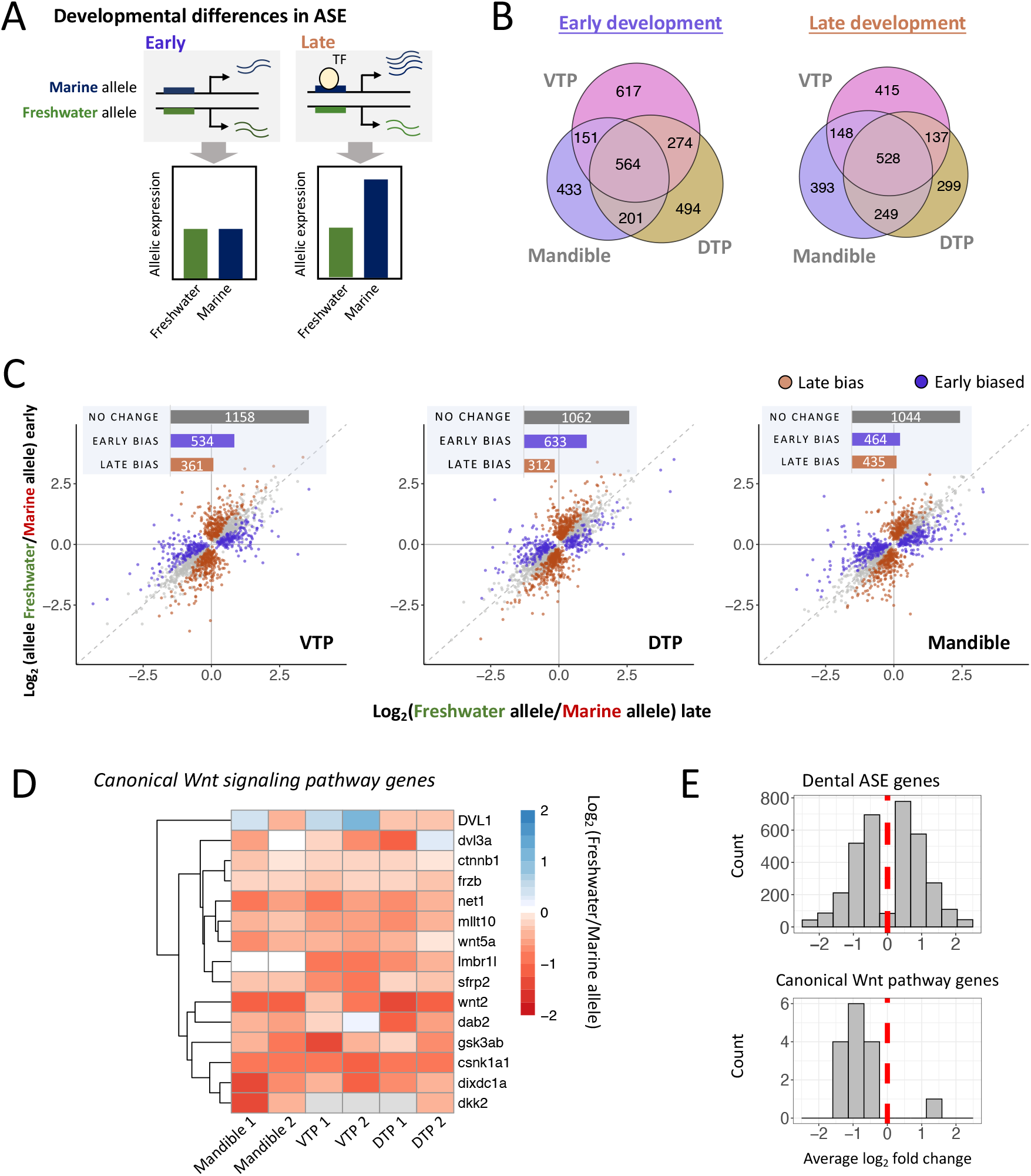
Developmental allele-specific expression in dental tissues. **A**. A schematic of differential ASE during development. In this example, sequence divergence between marine and freshwater sticklebacks at a *cis*-regulatory region results in allele-specific expression only in the presence of a context-specific transcription factor (“TF”, circle) expressed during late development. This results in differential allele-specific between developmental stages, shown in the bar plots. **B**. Venn diagram of ASE between dental tissues at the early (left) and late (right) developmental timepoint. **C**. Temporal differences in ASE during development in three dental tissues (left to right: ventral tooth plate, dorsal tooth plate, mandible). Genes with differential allele-specific between timepoints are colored based on magnitude of ASE in timepoint 1 vs. 2. Genes with greater differences in allelic expression in early development are shown in purple (“early bias”), genes with greater expression differences at the late timepoint are shown in terracotta (“late bias”). Gray points/bar (“no change”) indicate genes without evidence for significant differential allele-specific expression between timepoints. **D-E**. Genes involved in Canonical Wnt signaling show evidence of polygenic *cis*-regulatory evolution in dental tissues. Here we show genes from two Wnt signaling GO terms with biased directionality (Canonical Wnt signaling [GO:0060070]; Negative regulation of canonical Wnt signaling [GO:0090090]) (Table S7). In (**D**), the heatmap shows log2 fold changes for genes associated with canonical Wnt signaling and with ASE in at least one dental tissue. In (**E**), histograms of average ASE gene log2 fold changes from all dental genes (top) and the canonical Wnt signaling gene set (bottom). For each gene, log2 fold changes are averaged across any dental tissues in which ASE was identified.

Developmental stage was a major component of variation in allele-specific expression. Principal component analysis of marine-freshwater allelic log2 fold changes clustered tissues by timepoint, with late and early dental tissues forming separate clusters on PC2 (22% of the variance, Figure S5). PC analysis of allele-specific counts from individual tissues also clustered tissues based on developmental timepoint on either PC1 (mandible and DTP, 50% and 48% of variance, respectively) or PC2 (VTP, 33% of variance)(Figure S3).

In contrast to our comparison of more diverse tissues, dental ASE was often shared across tissues or developmental stages (Figures 3B, S6). Nearly 10% of genes with ASE in dental tissues (330/3471 genes) showed ASE in all three dental tissues and at both developmental timepoints. Overall, a greater proportion of genes with ASE were shared across dental tissues in late development compared to early development: 19% of ASE genes (564 genes) were shared across all three tissues in the early stage versus 26% (528 genes) in the late stage (Chi-square test, *P*=0.002). Examining stickleback orthologs of genes implicated in mammalian tooth development collected from the Bite-It and ToothCode databases (hereafter referred to as “BiteCode” genes [19]), we found that these genes were enriched for ASE (Fisher’s exact test, *P*=0.0046)(Figure S5B). BiteCode enrichment is consistent with the conservation of regulatory networks regulating dental development in mammals and fish [28,29].

Next, we characterized developmental differences in *cis*-regulatory divergence by comparing ASE between timepoints. Across developmental stages, divergent ASE can reflect the activity of temporally-specific genes controlled by divergent *cis*-regulatory elements between marine and freshwater fish (Figure 3A). Comparing the ratio of marine to freshwater allelic counts between early and late development in hybrids, we observed widespread differential allele-specific expression between the two developmental stages for each tissue, accounting for 37-43% of genes with ASE at either timepoint (Figure 3C)(Fisher’s exact tests, FDR<0.05; see Methods). The majority of differential ASE reflected ASE that was timepoint specific, meaning ASE was only observed at one developmental stage. However, roughly a quarter of differential ASE in each tissue was due to changes in the magnitude of ASE between timepoints.

More genes with differential ASE were found to have a larger *cis*-effect at the early stage than the late stage (i.e., |log2 fold change in early| > |log2 fold change in late|); Figure 3C), consistent with the greater proportion of genes with ASE at the early timepoint overall (Figure 1D). This result was surprising, as greater phenotypic divergence is observed between marine and freshwater fish in the pharyngeal tooth plates in late development [27]. Developmental differences were also typically tissue-specific: 67% of genes with developmental stage-bias ASE were unique to one tissue. Thus, *cis*-regulatory differences between marine and freshwater individuals are often specific to both tissue and tissue-developmental stage.

### Polygenic selection on cis-regulatory divergence between marine and freshwater sticklebacks

*Cis*-regulatory changes between marine and freshwater sticklebacks are potentially interesting for their role in local adaptation [8]. However, the majority of *cis*-regulatory changes are expected to be neutral. To test for selection on *cis*-regulatory changes between marine and freshwater fish, we employed a gene-set approach based on the sign test framework [30,31]. Under neutrality, QTLs for any given trait are expected to be unbiased with respect to their directionality, assuming these QTLs are independent (i.e. caused by different genetic variants) [32]. In a marine/freshwater genetic cross, each allele would be expected to be equally likely to increase the trait value if that trait is not under lineage-specific selection. Similarly, if a gene set associated with a biological function shows a significant directional bias in ASE (with more *cis*-changes acting in the same direction than expected), this suggests lineage-specific selection on the *cis*-regulation of this gene set [30,31]. Applying the sign test to GO gene sets in individual tissues and in the combined dental tissue set, we identified multiple gene sets with evidence for biased directionality (full list in Tables S5, S6).

In the combined dental tissue set, we found biased directionality for the GO terms “canonical Wnt signaling pathway” (Permutation based *P*-value = 0.0078), “embryonic viscerocranium morphogenesis” (*P* = 0.0093), and “Inflammatory response” (*P* = 0.0095). Wnt signaling plays a critical and evolutionarily conserved role in tooth and bone development [33,34] and genes in this pathway with ASE have been directly implicated in regulating dental development in other species (e.g., *Wnt5a, Sfrp2, Ctnnb1, Net1*)(Table S7). While the GO annotation term included both positive and negative regulators of Wnt signaling, the pathway “Negative regulation of canonical Wnt signaling” was also nominally significant for biased downregulation of freshwater alleles (7/7 genes, Fisher’s exact test, *P*=0.0048), suggestive of biased Wnt inhibition in marine fish (Figure 3D,E). Only three positive regulators of canonical Wnt signaling had ASE in dental tissues, precluding a separate statistical test of their directionality. As disruption or inhibition of canonical Wnt signaling results in arrested/aberrant tooth formation, selection on this pathway could potentially reflect selection for increased tooth number or related changes in feeding morphology in freshwater fish. Consistent with this, genes in the Wnt signaling pathway were previously shown to be upregulated in the VTP in PAXB freshwater compared to marine fish [19].

The GO term “embryonic viscerocranium morphogenesis”, which encompasses a set of genes involved in the generation and organization of the facial skeleton, also included ASE genes directly implicated in tooth and jaw formation (Table S8). For instance, *Dlx3b* and *Dlx1a*, genes encoding members of the Dlx family of homeodomain transcription factors [35], are involved in tooth and jaw patterning in mammals and fish [36,37]. Thus, biased directionality of this process category may reflect selection for morphological changes to the freshwater fish in the facial region related to feeding morphology.

We also found biased directionality for gene sets in individual tissues (Table S5). For instance, the GO category term “methyltransferase activity” (*P*=0.0018, 10/10 terms) showed biased upregulation of marine alleles in the eye and “endoplasmic reticulum” (*P*=0.0039, 38/50) showed biased upregulation of marine alleles in the flank skin. Since these gene sets are not yet associated with specific phenotypes, it is unclear what traits may have been impacted by their lineage-specific selection.

### Overlap between signatures of selection and genes with cis-regulatory divergence

If *cis*-regulatory changes underlie adaptive divergence between freshwater and marine forms, we may expect genes with ASE to fall within or near regions with signatures of selection. To test this hypothesis, we utilized a recent whole-genome analysis of differentiation between marine and freshwater populations from the northeast Pacific basin [9] (the source of the freshwater PAXB population studied here), where genomic regions of repeated marine-freshwater divergence were identified through marine-freshwater cluster separation scores (CSS). A CSS score quantifies average marine-freshwater genetic distance after subtracting the genetic distance found within each ecotype for a genomic window [8,9].

We asked whether genes with ASE co-localized with genomic regions with greater evidence for marine-freshwater divergence (i.e., greater CSS Z-scores). As power to detect ASE is related to the number of variant sites, we compared median CSS Z-scores between ASE and background genes with similar SNP densities (see Methods, Figure S7). Genes with ASE were associated with greater Z-scores per SNP density bin (Figure 4A,B; Permutation *P*<0.0001), indicating an enrichment of ASE genes in genomic regions with greater evidence for marine-freshwater divergence. This pattern is consistent with the hypothesis that repeated marine-freshwater divergence may often involve changes in gene regulation [8,18,18].

**Figure 4.**
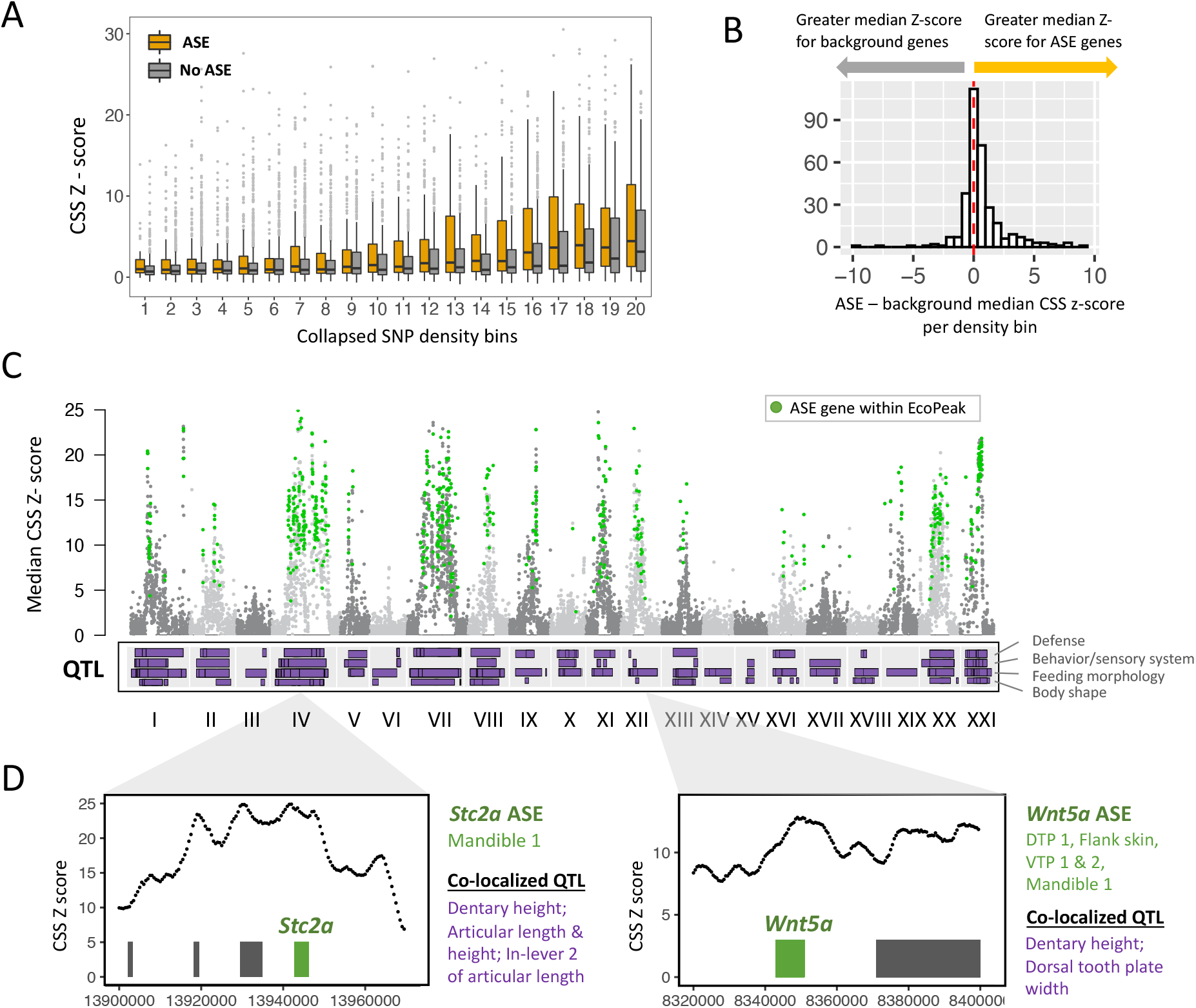
ASE genes are associated with regions of repeated marine-freshwater divergence. **A**. Average marine-freshwater cluster separation score (CSS) Z-scores for ASE genes and background genes binned by SNP density. Here, genes are separated into 20 density bins for visualization, with higher numbers corresponding to greater SNP density. **B**. ASE genes have higher average CSS Z-scores than background genes. The histogram shows median Z scores for ASE genes minus background genes for each SNP density bin. More density bins show positive values, indicating higher average Z-scores for ASE genes overall (Permutation *P*<0.0001). **C**. Manhattan plot of median gene CSS Z score vs. chromosome position. Highlighted in green are genes with ASE that overlap significant regions of recurrent marine-freshwater divergence in the northeast Pacific basin (“EcoPeaks”). Below we show locations of quantitative trait loci (QTLs) identified in previous genetic crosses between PAXB and marine fish. QTL are divided into three broad categories (from top to bottom: defense, behavior and sensory system, feeding morphology, and body shape). **D**. Two candidate ASE genes within regions of marine-freshwater divergence. ASE was observed for *Wnt5a* and *Stc2a* in one or more dental tissues and these genes co-localize with QTL related to feeding morphology. Bars in the panel indicate genes within these regions, with candidate genes *Wnt5a* and *Stc2a* highlighted in green. Tissue(s) in which ASE was identified (green) and relevant overlapping QTL (purple) are listed to the right of each gene panel.

Regions with significant CSS scores (EcoPeaks) overlapped 611 ASE genes (13.8% of ASE genes overall; 1.9-fold enrichment, Permutation test *P*<0.001)(Figure 4C, Table S9). Genes with evidence for ASE heterogeneity between tissues were enriched within EcoPeaks compared to all genes with evidence for ASE (Fisher’s exact test, *P*=0.01), as were genes with evidence for tissue-specific ASE (Fisher’s exact test, *P*=0.018).

Marine-freshwater EcoPeaks are clustered throughout the genome, which is thought to reflect selection on linked “supergene” complexes affecting multiple traits [9,38]. We also find that genes with ASE are enriched on particular chromosomes (Figures S8, S9; see Methods). EcoPeaks and QTL associated with phenotypic divergence are particularly concentrated on ChrIV and this chromosome also harbored the highest proportion of ASE genes over background (Permutation *P*<0.001) as well as a quarter of EcoPeak ASE genes (154 genes). A more modest enrichment of ASE genes was also found for chrXXI (*P*=0.034) and chrXI (*P*=0.033), which have been shown to harbor inversions between marine and freshwater fish [8]. We identified 56 and 48 ASE genes within EcoPeaks on these chromosomes, respectively.

To identify potential candidate genes for marine-freshwater divergence, we overlapped ASE genes identified in marine-freshwater peaks with QTL for dental and skeletal traits [6,22,39](Figure 4C,D). QTL for variation in dental traits between PAXB freshwater and marine fish (e.g., VTP or DTP tooth plate size and shape, tooth number, and jaw size and shape) overlapped 401 genes with ASE in relevant tissues (File S1). A small subset of these have previously been implicated in dental or craniofacial morphology in other species (Table 1), including several genes involved in the Wnt signaling pathway identified in the sign test (e.g., *Wnt5a, Sfrp2, Ctnnb1, Net1*).

**Table 1.**
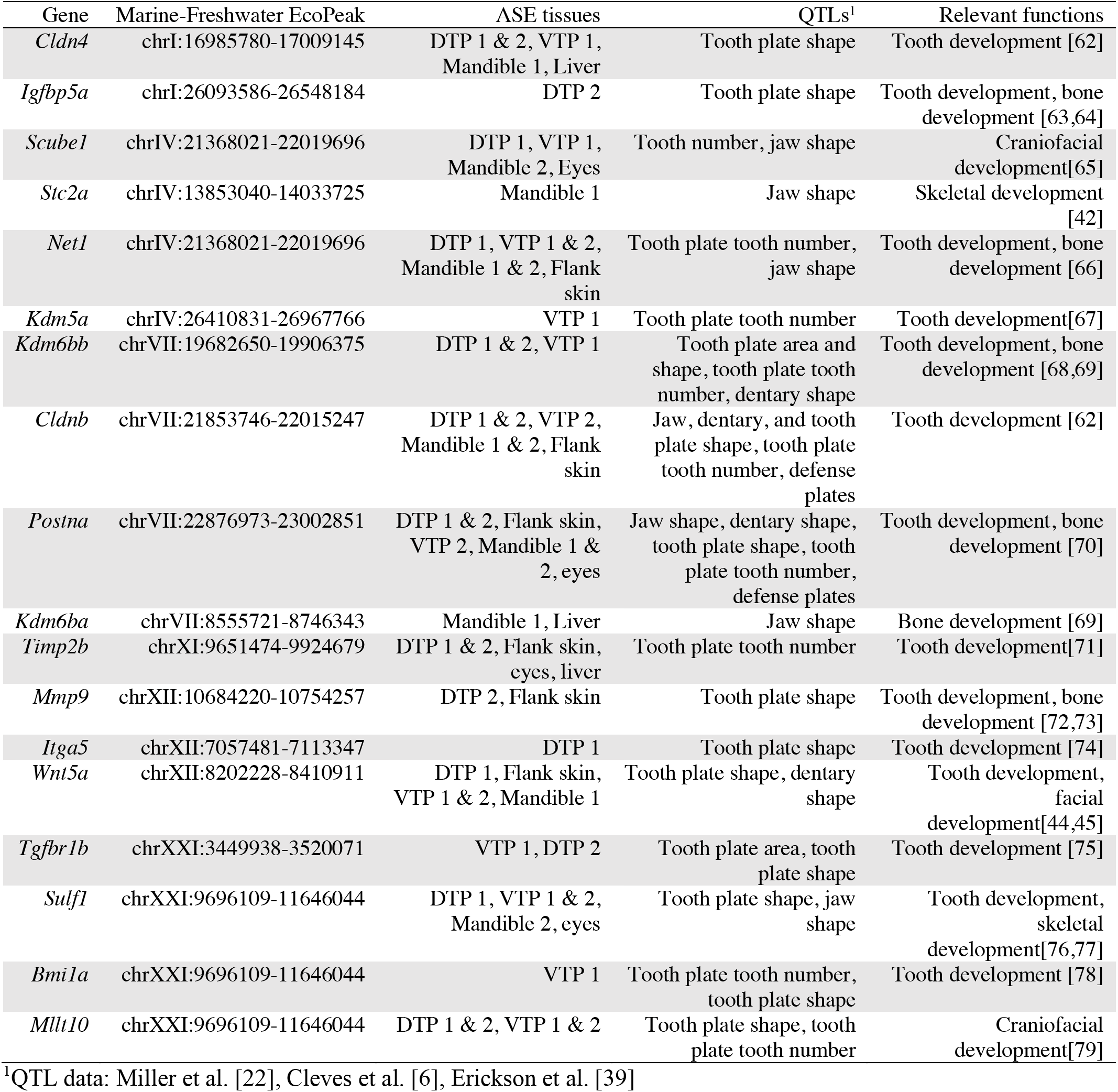
Candidate ASE genes within differentiated regions with overlapping QTL.

## Conclusions

Changes in gene expression regulation are thought to play a major role in evolutionary adaptation. Here, we surveyed allele-specific expression across tissues and developmental stages to understand the landscape of *cis*-regulatory divergence between marine and freshwater sticklebacks. We identified widespread ASE that was largely heterogeneous between tissue types and developmental stages. For a subset of these *cis*-regulatory changes, we found evidence for polygenic selection on particular processes/pathways with a sign test. Finally, we demonstrated that *cis*-regulatory changes are often associated with regions of marine-freshwater divergence, further supporting the role of *cis*-regulatory differences in adaptive evolution in sticklebacks [8,9].

Our results indicate the *cis*-regulatory divergence between marine and freshwater fish is often specific to an individual tissue or developmental stage. Gene expression differences that are spatially or temporally restricted may be important in the process of adaptation to new environments. Through context-specific expression regulation, *cis*-regulatory mutations can avoid negative pleiotropy associated with global changes in expression or protein structure. Thus, it is possible that *cis*-regulatory variation that introduces discrete changes in gene expression may be favored during adaptation. Interestingly, we found that genes with evidence for tissue-specific ASE in particular were enriched in regions of recurrent marine-freshwater divergence. Tissue- or context-specific *cis*-regulatory differences have previously been shown to underlie adaptive traits in sticklebacks [4,14] and other systems[40]. The tissue- and developmental-specificity of *cis*-regulatory changes we identified between marine and freshwater sticklebacks highlights the utility of studying gene regulation across multiple tissues and contexts in understanding regulatory adaptation.

Genes with ASE in regions of repeated marine-freshwater divergence may be interesting candidates for adaptive phenotypic differences between ecotypes. *Cis*-regulatory changes have been found to underlie a number of phenotypic differences between marine and freshwater forms. For example, *cis*-regulatory changes at *Bmp6* are associated with evolved tooth gain[6,15], and *cis*-regulatory changes at *Eda* and *GDF6* have been implicated in skeletal differences between marine and freshwater fish[7,41]. While we did not have sufficient expression to examine ASE at these genes specifically (e.g., *Eda, GDF6, Bmp6*) in our dataset, we identified a number of potentially interesting candidate genes within differentiated genomic regions with ASE. For example, *cis*-regulatory variation at *Stanniocalcin2a (Stc2a*) was recently associated with changes in spine length in freshwater sticklebacks [16]. We found that *Stc2a* also showed ASE in the early mandible timepoint. *Stc2a* falls within a marine-freshwater divergent region on ChrIV that overlaps several QTL, including the QTL with the largest effect on dentary size in crosses between PAXB freshwater fish and marine fish [22]. In mice, *Stc2a* modulates bone size and growth and overexpression results in smaller mandibles[42,43], making this gene an exciting candidate for divergent jaw morphology between marine and freshwater benthic fish. Our results also highlighted Wnt signaling genes as potential candidates for divergence in feeding morphology. A sign test indicated evidence for lineage-specific selection on *cis*-regulatory alleles involved in Wnt signaling, and several of these genes were also found within QTL/EcoPeaks and involved in tooth development or craniofacial morphology (see Table S1). For example, *Wnt5a*, associated with QTL for tooth plate and dentary shape, plays an important role in facial and tooth development in mammals [44–46]. Our results establish the landscape of stickleback *cis*-regulatory divergence across tissues and developmental stages; we look forward to future studies that elucidate the roles that specific ASE genes have played in stickleback adaptation.

## Methods

### Stickleback husbandry

All animal work was approved by UC Berkeley IACUC protocol AUP-2015–01-7117. Fish were raised in aquaria at 18°C in brackish water (3.5g/L Instant Ocean salt, 0.217mL/L 10% sodium bicarbonate) with 8 hours of light per day. Fry (SL < 10mm) were fed live Artemia, early juveniles (SL ~10-29 mm) were fed live Artemia and frozen Daphnia. Fish above ~20mm were fed frozen bloodworms and Mysis shrimp. To generate F1 hybrids, a freshwater Paxton Benthic (Paxton Lake, Canada) strain male was crossed with a marine Rabbit Slough (Alaska) strain female. Individuals from these lineages have been maintained in the lab for >10 generations. The resulting full-sibling fish were raised together in a common dish or tank until sample collection. Female F1 hybrids were selected for dissection at two timepoints (15-20 mm SL and 35mm SL). Fish were euthanized individually via immersion in 250 mg/L MS-222. Tissue samples for RNA-seq were immediately dissected on an ice-cold tray. Brain samples included all bilateral brain regions from the olfactory bulb to the brain stem. Liver samples were derived from the anteriormost lobe of the fish liver. Eye samples encompassed the entirety of the left eye of each fish, including the majority of the optic nerve. Flank skin samples were taken by removing the majority of the skin covering the left side of each fish, capturing a region that would normally be covered by lateral armor plates in adulthood (anteriormost boundary at the level of the 1^st^ dorsal spine, where the anteriormost armor plates had begun ossification at the time of dissection, posterior boundary at the back of the dorsal fin where armor plates were not yet ossified). Dorsal pharyngeal tooth plate samples included left and right DTP1 and DTP2, as well as underlying epibranchial bones and surrounding soft tissues and teeth. Ventral pharyngeal tooth plate samples included left and right ceratobranchial 5 and surrounding soft tissues and teeth. The mandible consisted of the dentary bone and lower lip, and all associated soft tissues and teeth. Samples were placed into 50 ul of TRIzol (Invitrogen), briefly agitated by shaking, and incubated on ice for 10 minutes. All samples from each timepoint were all prepared on the same day.

### RNA extraction, library preparation, and sequencing

Dissected tissues were kept in TRI reagent and stored in −80°C prior to RNA-extraction. Total RNA extraction was performed as described previously [19]. Total RNA was quantified by Qubit Fluorometer, and quality was checked by Agilent Bioanalyzer. Libraries were constructed with New England Biolabs NEBNext Poly(A) mRNA Magnetic Isolation Module (E7490S), NEBNext Ultra II Directional RNA Library Prep Kit (E7765S) and NEBNext Multiplex Oligos for Illumina (96 Unique Dual Index Primer Pairs, E6440S) following the manufacturer’s instructions. Library quality was analyzed on an Agilent Bioanalyzer (Table S1). Libraries were pooled and sequenced on an Illumina HiSeq platform (2×150 bp reads). We obtained a total of 1,439,700,457 reads across 20 samples (10 tissues x 2 replicates) (Table S1).

### Whole-genome re-sequencing of PAXB

To phase RNA-seq reads, whole-genome resequencing was performed on the PAXB parent. DNA was extracted from fin tissue. Library preparation and sequencing were performed by Admera Health (South Plainfield, NJ). Libraries were sequenced on an Illumina HiSeqX platform (2×150 bp reads) to a depth of ~30X (Figure S1). Coverage per site was calculated with Samtools depth [47] based on reads aligned to the reference genome (described below).

### Read mapping and SNP calling

RNA-seq read quality was assessed using FastQC. Reads were trimmed for adaptor sequences with Trimmomatic[48] and then mapped to the stickleback reference genome [49]. F1 hybrid RNA-seq reads were mapped to the stickleback reference genome with STAR v2.7 [50]. Genomic reads from PAXB were mapped with bowtie2 v2.3.4 (argument: --very-sensitive) [51].

SNP calling was then performed with the Genome Analysis Tool Kit (GATK)[52]. Duplicates were marked with the Picard tool MarkDuplicates. Read groups were added with AddOrReplaceReadGroups. For RNAseq reads, we used GATK tool SplitNCigarReads to split reads that contain Ns in their cigar string (e.g., spanning splice events). GATK HaplotypeCaller and GenotypeGVCFs were used for joint genotyping. SNP calls were subsequently filtered for low quality calls with VariantFiltration (QD < 2.0; QUAL < 30.0; FS > 200; ReadPosRankSum < −20.0).

To assign allele-specific reads to the parent of origin (i.e., “freshwater” parent vs. “marine” parent), we retained only variants where the PAXB parent was homozygous. Heterozygous sites for each F1 individual were used for separating allele-specific RNA-seq reads into freshwater and marine pools, as described below.

### Identifying allele-specific expression

To identify allele-specific expression (ASE), reads from each library were then mapped again with STAR, implementing the WASP filter based on heterozygous calls [53]. WASP reduces mapping bias by identifying reads containing SNPs, simulating reads with alternative alleles at that locus, re-mapping these reads to the reference, and then flagging reads that do not map to the same location. Reads that do not map to the same location were discarded[53]. Parental origin for each allele was assigned based on PAXB (freshwater parent, see above). Reads were counted over marine-freshwater variants with ASEReadCounter [52] for individual heterozygous sites. To mitigate the effects of SNP calling errors and read mapping bias, we removed heterozygous sites with: 1) large ratio differences indicative of mapping bias (log2 fold changes of allelic counts > 10), or 2) no reads mapped to one of the parental alleles. Mapping was then repeated a second time based on the updated list of heterozygous sites. Analysis of ASE ratios in each library centered around a log2 ratio of zero, indicating approximately equal mapping to both parental alleles.

Gene-wise estimates of allele-specific expression were quantified by counting allele-specific reads overlapping exons using HTSeq [54] based on Ensembl annotations (BROAD S1) [8], with coordinates converted by LiftOver to the v4 stickleback assembly (https://stickleback.genetics.uga.edu/downloadData/)[49]. Total counts per parental allele per tissue are available in Table S2. Across tissues, we did not observe a consistent bias towards either of the parental alleles. To examine transcriptome-wide patterns of expression, we transformed expression values (allele-specific and total counts) using variance stabilizing transformation and assessed transcriptome-wide expression patterns via principal components analysis (PCA)(Figures 1B, S2).

DESeq2 [55] was used to identify ASE using the individual as a blocking factor and allele-specific expression (“marine” vs. “freshwater” allele) as the variable of interest (Wald test). As read counts from “marine” and “freshwater” alleles come from the same sequencing library, library size factor normalization was disabled by setting SizeFactors = 1. *P*-values were adjusted using the Benjamini-Hochberg method in DESeq2 for multiple comparisons. Genes were examined at FDR<0.05 and FDR<0.1 (Table S3). Comparing genes with ASE in VTP from the late timepoint with the results of Hart et al. [19], which also tested for ASE in crosses between PAXB and RABS in the VTP, we found highly significant overlap (Fisher’s exact test, *P*=8.83×10^-292^). Fifty-one percent of ASE genes identified here were also identified in the previous analysis. Additionally, log2 fold changes of genes with ASE were found to be correlated (Pearson’s Correlation, *r* =0.64, *P*=2.47×10^-69^).

Developmental stage differences in ASE were identified by comparing reads mapping to freshwater vs. marine alleles at both timepoints, summed across the two replicates. We compared marine and freshwater allelic ratios for genes with evidence of ASE in at least one of the two developmental stages with a Fisher’s exact test. Resulting *P*-values were corrected using the Benjamini-Hochberg method.

### Assessing heterogeneity in allele-specific expression across tissues

To identify heterogeneity in allele-specific expression across tissues, we adopted a Bayesian model comparison framework from Pirinen et al. [23]. In this approach, tissues are classified as no ASE 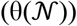, strong ASE 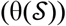, or moderate ASE 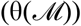 based on freshwater and marine allelic counts summed across replicates per gene under a grouped tissue model [23]. Tissues are further classified as showing ASE heterogeneity if one tissue showed evidence for either strong or moderate ASE and at least one other tissue did not show ASE (HET0) or when all tissues showed some evidence for ASE but the magnitude differed (HET1). Finally, we consider a sub-state of ASE heterogeneity tissue-specificity, where ASE state (i.e., moderate, strong ASE, or no ASE) is observed in only one tissue [23].

The following priors were selected to describe groups:

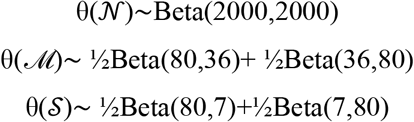

Densities of the prior distributions for the proportion of allelic counts are found ins Figure S9. Parameters for Beta distributions were chosen to clearly separate the three groups from each other to allow the classification of tissues to a particular group, following Pirinen et al. [23]. The “No ASE” condition dominates the region around 0.5 (0.47,0.53), allowing for some deviation for technical bias or noise [23,56]. Strong ASE dominates at extreme frequencies ([0.85,0.96],[0.3,0.15]) and moderate ASE dominants between these two groups (Figure S9). Genes expressed in at least two tissues at a minimum depth of 10 reads/allele were included in the analysis (15,477 genes). For each gene, we excluded tissues for which coverage was low (less than or equal to 10 reads per allele).

### Sign test on ASE

To search for selection on cis-regulatory variation, we applied a sign test based on the directionality of ASE in a gene set [30,31]. Gene Ontology (GO) categories for zebrafish were obtained from ZFIN (https://zfin.org/downloads)[57] and mapped to stickleback orthologs based on Ensembl ortholog annotations. Genes with evidence for *cis*-regulatory divergence were divided into categories based on the upregulated allele (freshwater vs. marine). We excluded GO categories with fewer than 10 members in a test set with ASE. As test sets contain different proportions of upregulated marine vs. freshwater alleles, we tested for lineage-specific bias in each test set with a Fisher’s exact test.

Because many GO categories were tested, we determined the probability of an enrichment by permuting gene category assignments, as described previously [30,31,58]. Gene assignments were shuffled and the test was repeated 10,000 times. Permutation-based *P*-values were determined by asking how often a result of equal or greater significance would be observed in permuted datasets [30,31,58]. Tests were performed on individual tissues and on the dental tissues together, as many genes with ASE are shared across these tissues. In the grouped tissue analysis, we looked for biased directionality across all genes with ASE in a tissue group. In the event that signs differ between tissues (i.e., freshwater allele is upregulated in tissue #1, marine allele is upregulated in tissue #2), the gene is discarded from the analysis. Changes in directionality across tissues was only seen for one gene associated with a significant GO term (dental tissue: “inflammatory response”). To ensure that biased directionality in our group analyses were robust to tissue-specific patterns, we performed a second test where we combined *P*-values for a tissue group from individual tissues with Fisher’s method, as in [31]. We performed a Fisher’s exact test for each category as described above for individual tissues. *P*-values for GO categories that are represented across all tested groups were then combined using the R package metap. We report on GO terms significant in both approaches, as these represent cases of biased directionality across tissue groups and robust to individual tissue patterns. Combined *P*-values are reported in Table S6.

### Identifying overlap with EcoPeaks

Data from Kingman et al. [9] was downloaded from the UCSC Genome Browser Table Browser ([22]: https://sbwdev.stanford.edu/kingsleyAssemblyHub/hub.txt). Intervals were associated with overlapping genes using bedtools [59]. As power to detect ASE is related to the density of informative sites, we calculated SNP density per gene as the number of informative heterozygous sites divided by transcript length, based on BROAD S1 gene annotations [9]. To determine whether genes with ASE had higher average Z-scores than background genes, while controlling for the effect of SNP density on our power to identify ASE, we grouped genes with similar SNP densities into bins based on the distribution of SNP density values. We excluded bins for which there were fewer than five genes in each category (ASE, no ASE) so as not to skew results based on bins with few observations (330 bins, average of 23 genes per bin)(Figure S8A). For each bin, we calculated the median Z-score for genes with ASE and for background genes (Figure S8B). We compared this result to a permuted dataset. Within a density bin, we shuffled gene category assignments and again calculated the median Z-score for each category in each bin. We repeated this 10,000 times. To obtain a permutation-based *p*-value, we compared how often the median difference in category Z-scores was as extreme or more extreme than in empirical data. This result was robust to varying bin sizes (Table S10).

To ask whether ASE genes were enriched on particular chromosomes, as with QTLs and EcoPeaks [9], we performed a resampling test to account for differences in SNP density between genes. We sampled random sets of genes (equal to the number of total ASE genes) with SNP densities matched to the ASE gene set (1000 times). The number of genes associated with each chromosome were counted for each permuted gene set and compared to the empirical data. *P*-values were calculated based on how often an equal or more extreme result was observed for permuted gene sets (Figure S9).

### Gene annotations and QTL overlap

Stickleback genes were annotated to zebrafish and mouse orthologs based on Ensembl ortholog annotations. BiteCode genes were annotated as in Hart et al. [19], from the BiteIt database (http://bite-it.helsinki.fi/) and ToothCODE database (http://compbio.med.harvard.edu/ToothCODE/). Phenotype annotations for zebrafish were downloaded from ZFIN (https://zfin.org/downloads), mouse mutant phenotypes were downloaded from Ensembl and the Mouse Genome Database[60].

QTL coordinates for overlap are based on genomic coordinates in Marques and Peichel [61]. For overlap with ASE genes, we focused on QTL mapping studies utilizing crosses between PAXB freshwater and marine individuals. Dental QTL for Table S1 and File S1 were obtained from three studies [6,22,39]. Coordinates were converted by LiftOver to the v4 stickleback assembly for overlap with EcoPeaks. Genes of interest for Table 1 were identified based on intersections between these genes (ASE/EcoPeak/QTL) and phenotype/Gene Ontogony annotations or the BiteCode gene list. A full list of gene overlaps is available in File S1.

## Data Availability

All sequence data generated in this study have been deposited to the National Center for Biotechnology Information Sequence Read Archive as a BioProject (SUB12080818). Supplemental datasets are available in File S1. Scripts associated with this manuscript are available on GitHub (https://github.com/katyamack-hub/SticklebackASE).

## Supplemental Material

Supplement Figures and Tables

Figues S1-S10

Tables S1-S10

Supplemental references

File S1: Supplemental gene lists

