## Supplemental Figures and Tables for "Evolution of spatial and temporal *cis*-regulatory divergence between marine and freshwater sticklebacks"

**Supplemental Material**

**Supplemental Figures**

**Figure S1**. Coverage of PAXB genome re-sequencing.

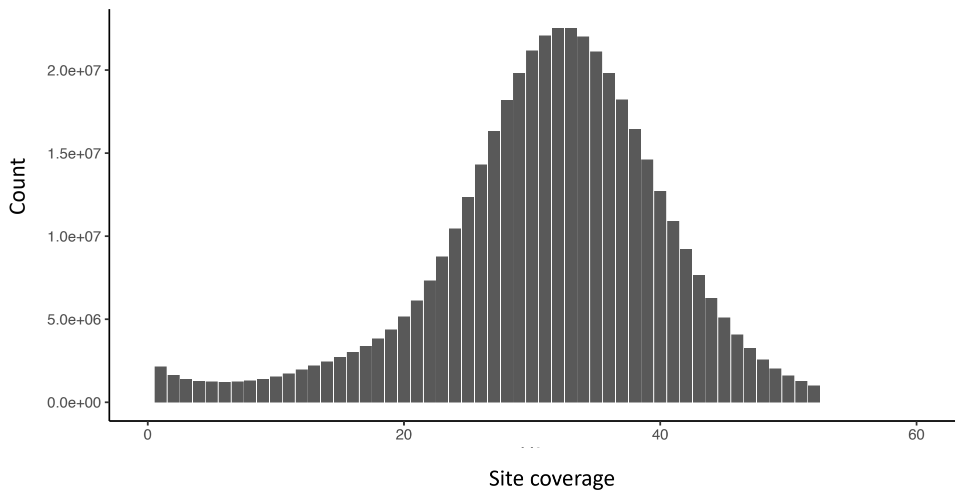

**Figure S2.** PCA of total RNAseq reads across tissues of marine-freshwater F1s.

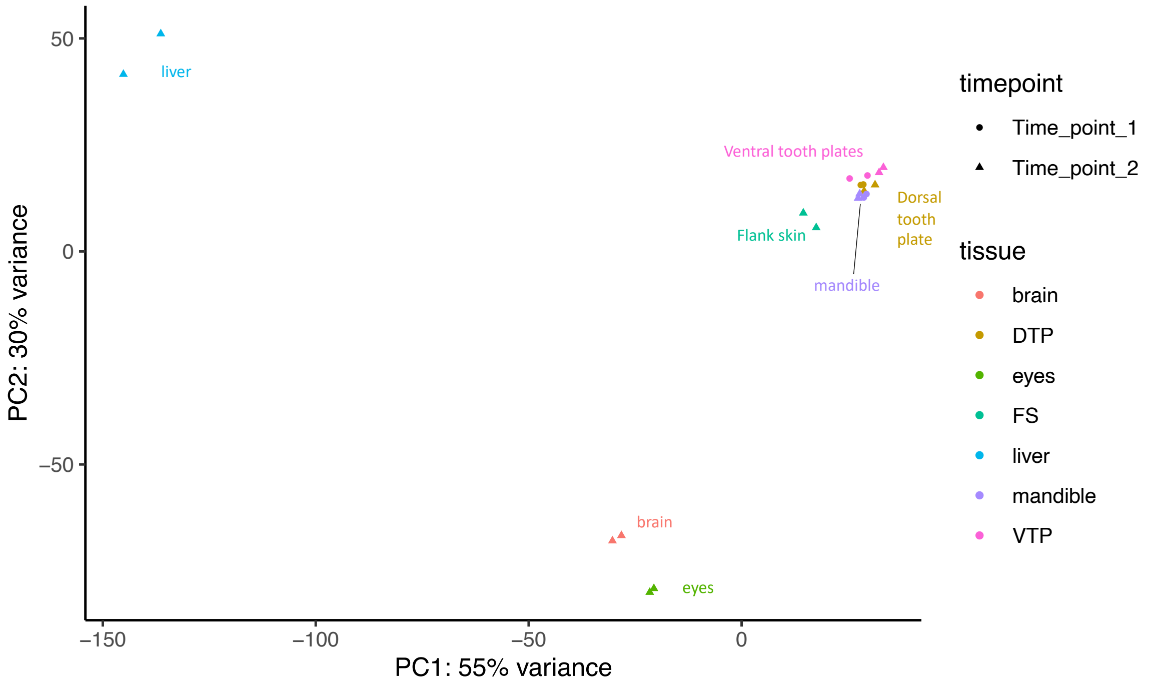

**Figure S3**: PCA of dental tissues from two developmental stages. Shape indicates freshwater (triangle) vs. marine allele (circle) and color indicates developmental stage (time 1 = red, time 2 = teal).

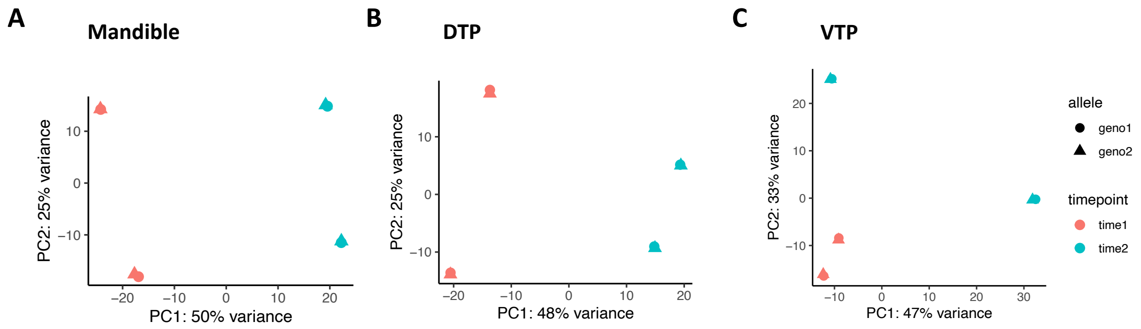

**Figure S4.** Total allele-specific reads per tissue vs. number of genes with evidence for ASE in a tissue (at FDR<0.05)**.** Each dot represents a tissue. Allele-specific read depth was not correlated with number of ASE genes identified in a tissue (Spearman's rank correlation, *p*=0.56, *rho*=-0.24).

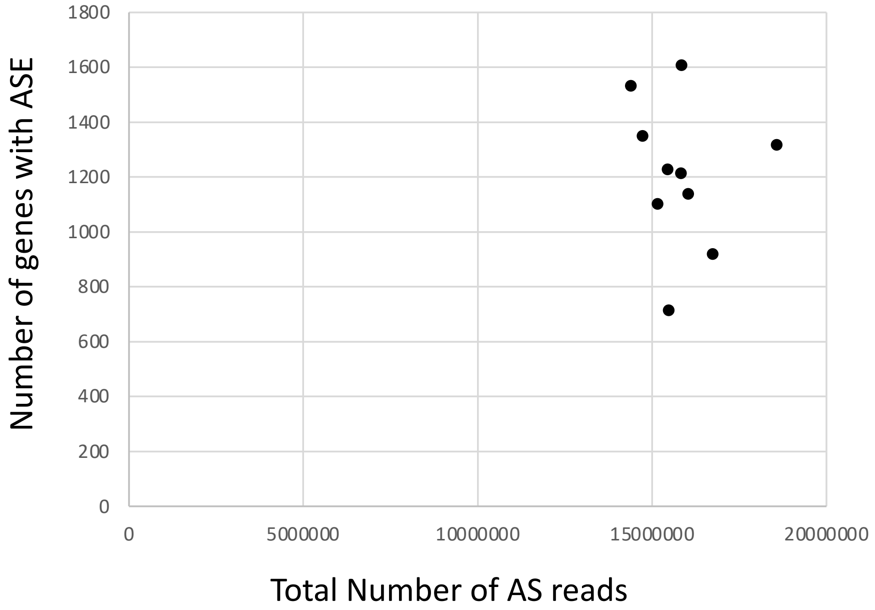

**Figure S5. A**. PCA of average log2 fold change (marine/freshwater allele) of dental tissues. **B**. BiteCode genes with ASE. Genes are colored by their log2 fold change. Asterisks indicate significance level *FDR<0.05, **FDR<0.01, ***FDR<0.001.

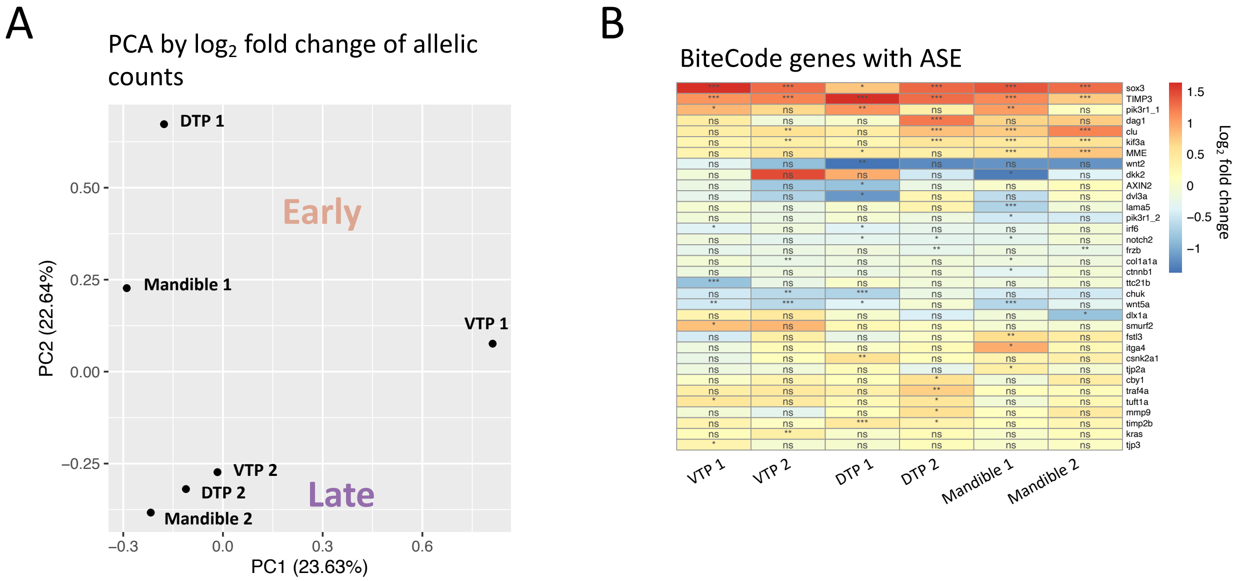

**Figure S6**. UpSet plot of overlap of ASE across dental tissues at two developmental stages.

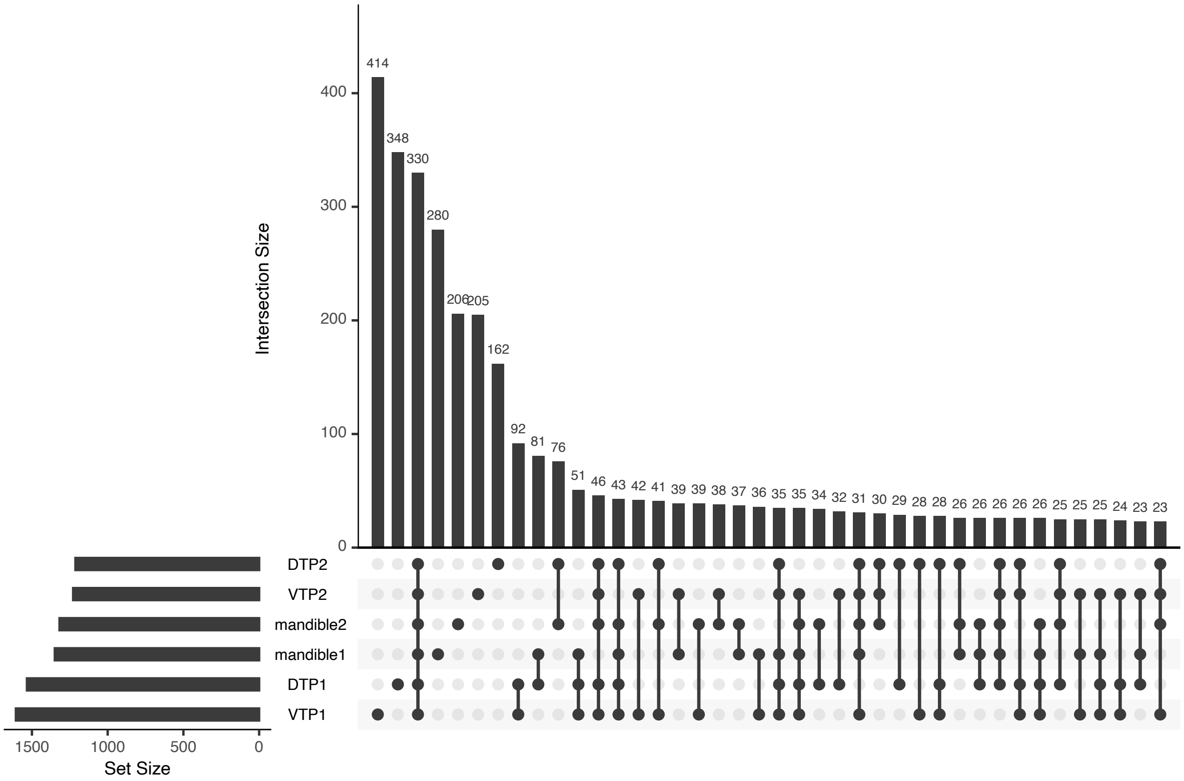

**Figure S7. A**. Density plots of average SNP density in bins for ASE genes (right) and background genes (left). A similar density distribution is seen for ASE vs. background for bins. **B.** Histogram of median Z-scores of ASE genes and background genes for bins (bars are interleaved for visualization). **C.** Gene median Z scores vs. bin. Each point is a gene.

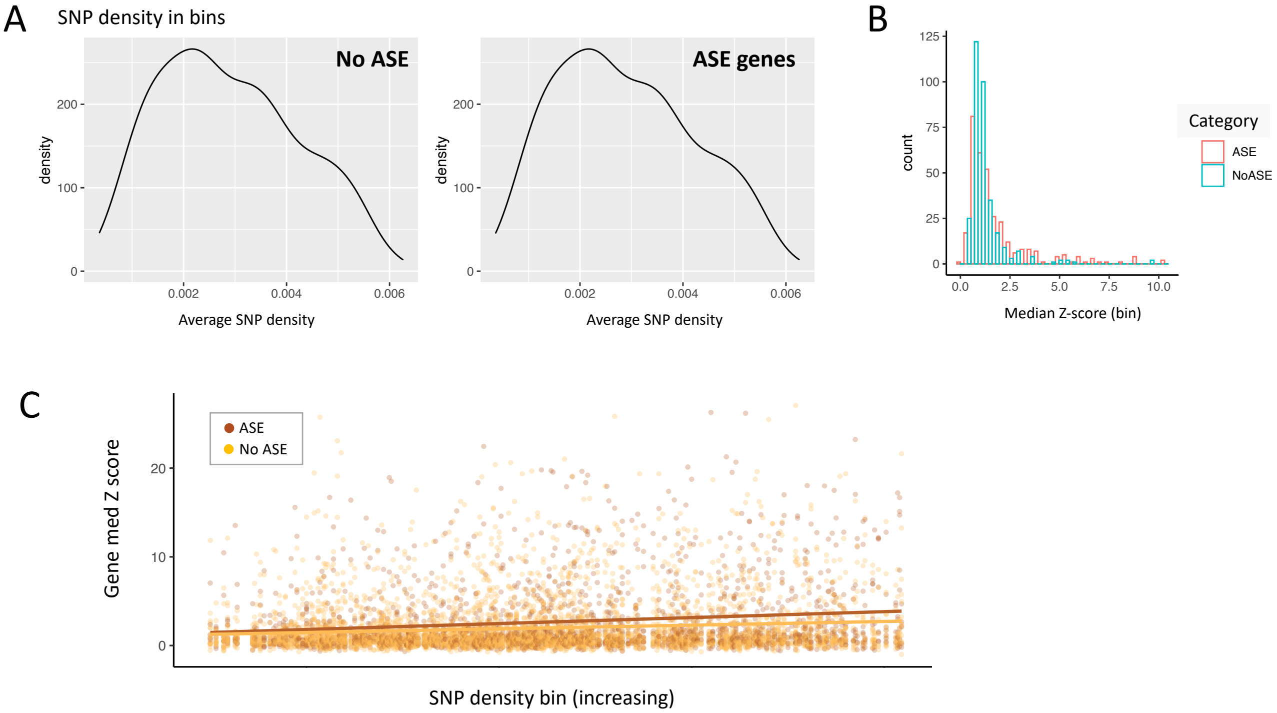

**Figure S8.** Proportion of gene on each chromosome with ASE for each tissue (number of genes with ASE/background on a chromosome)

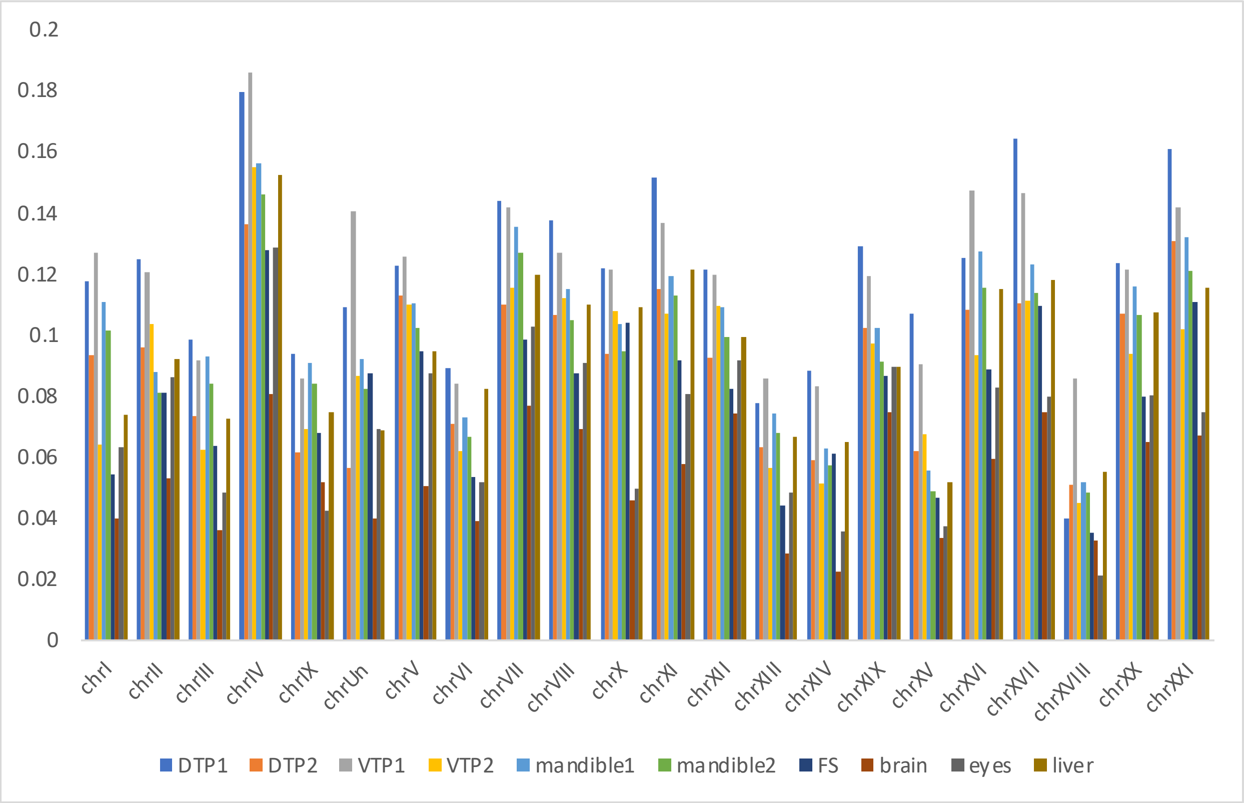

**Figure S9.** Chromosomes with an enrichment of ASE over background expression (*P*-values reported based on permutation).

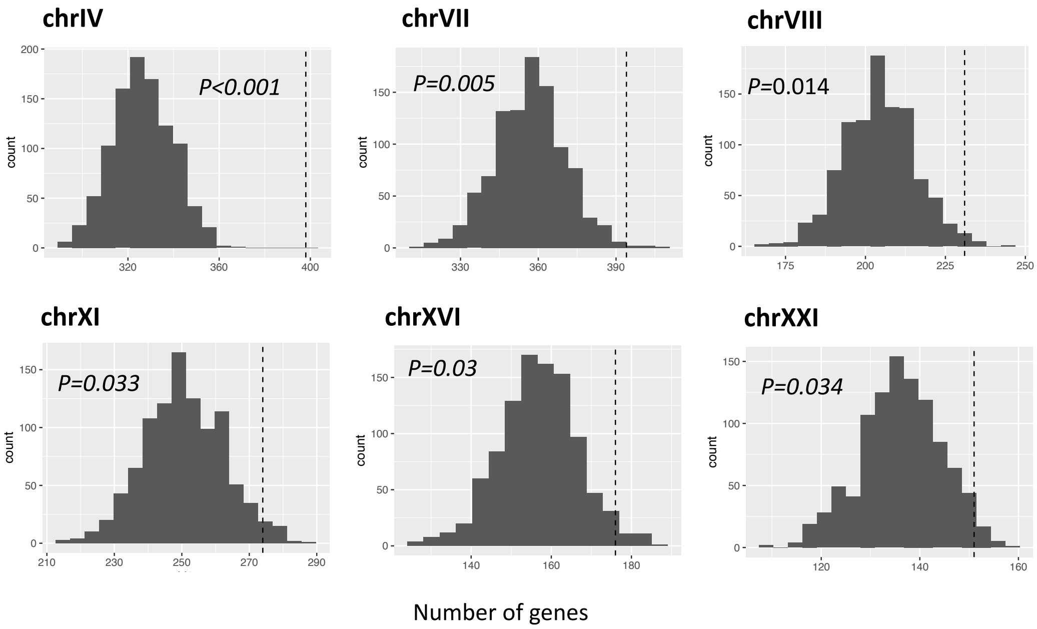

**Figure S10.** Densities of the prior distributions for the proportion of the freshwater allele. Parameters for Beta distributions were chosen to clearly separate the three groups from each other to allow the classification of tissues to a particular group (no ASE ($\theta(\mathcal{N})$), strong ASE ($\theta(\mathcal{S})$), or moderate ASE ($\theta(\mathcal{M})$)), following Pirinen et al. [1].

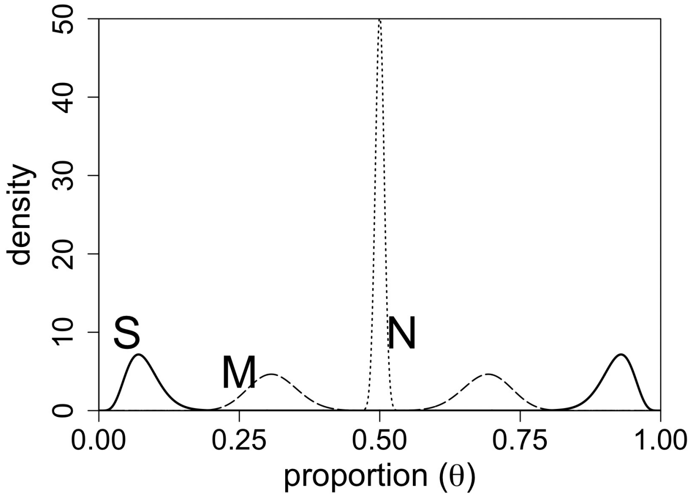

**Table S1.** RNAseq library sample information

| Sample_ID | Tissue | Timepoint | RIN score | Raw reads | Mapped Reads (total) |
|---|---|---|---|---|---|
| 18146FL-37-01-01_S74 | brain | Time point-2 | 9.8 | 83303949 | 75363055 |
| 18146FL-37-01-02_S75 | brain | Time point-2 | 10 | 70737373 | 63643543 |
| 18146FL-37-01-05_S78 | DTP | Time point-1 | 7.7 | 66748054 | 57054598 |
| 18146FL-37-01-06_S79 | DTP | Time point-1 | 8 | 66666757 | 58252092 |
| 18146FL-37-01-03_S76 | DTP | Time point-2 | 9.5 | 82208758 | 73258066 |
| 18146FL-37-01-04_S77 | DTP | Time point-2 | 8.9 | 61076908 | 54068495 |
| 18146FL-37-01-07_S80 | eyes | Time point-2 | 9 | 66645858 | 59796162 |
| 18146FL-37-01-08_S81 | eyes | Time point-2 | 10 | 91056685 | 82687823 |
| 18146FL-37-01-09_S82 | FS | Time point-2 | 9.8 | 62002949 | 54244891 |
| 18146FL-37-01-10_S83 | FS | Time point-2 | 10 | 69199627 | 61208309 |
| 18146FL-37-01-11_S84 | liver | Time point-2 | 10 | 66280457 | 60093436 |
| 18146FL-37-01-12_S85 | liver | Time point-2 | 9.8 | 66475414 | 59547137 |
| 18146FL-37-01-13_S86 | mandible | Time point-1 | 7.4 | 63449422 | 55541248 |
| 18146FL-37-01-14_S87 | mandible | Time point-1 | 8.7 | 65477982 | 57454455 |
| 18146FL-37-01-15_S88 | mandible | Time point-2 | 10 | 99370852 | 87241465 |
| 18146FL-37-01-16_S89 | mandible | Time point-2 | 10 | 65309621 | 58272877 |
| 18146FL-37-01-17_S90 | VTP | Time point-1 | 10 | 71780930 | 62100619 |
| 18146FL-37-01-19_S92 | VTP | Time point-2 | 7.9 | 92388481 | 81080118 |
| 18146FL-37-01-20_S93 | VTP | Time point-2 | 7.9 | 52927806 | 46928285 |
| 18146FL-37-01-18_S91 | VTP | Time point-1 | 8.1 | 76592574 | 64309864 |

**Table S2**. Allele-specific mapping depth per sample.

| Individual | Freshwater allele | Marine allele |
|---|---|---|
| DTP_1, replicate 1 | 3745530 | 3685882 |
| DTP_1, replicate 2 | 3473514 | 3474894 |
| DTP_2, replicate 1 | 4692653 | 4688887 |
| DTP_2, replicate 2 | 3211629 | 3216112 |
| FS_2, replicate 1 | 3686618 | 3524350 |
| FS_2, replicate 2 | 4003882 | 3930078 |
| VTP_1, replicate 1 | 3956071 | 3901128 |
| VTP_1, replicate 2 | 3984927 | 3982959 |
| VTP_2, replicate 1 | 4836454 | 4777676 |
| VTP_2, replicate 2 | 2917434 | 2897866 |
| brain_2, replicate 1 | 4238704 | 4267511 |
| brain_2, replicate 2 | 3452370 | 3501538 |
| eyes_2, replicate 1 | 4751740 | 4821184 |
| eyes_2, replicate 2 | 3542170 | 3604704 |
| liver_2, replicate 1 | 4144471 | 4127935 |
| liver_2, replicate 2 | 3917175 | 3829309 |
| mandible_1, replicate 1 | 3590991 | 3473673 |
| mandible_1, replicate 2 | 3876991 | 3769969 |
| mandible_2, replicate 1 | 5617035 | 5452744 |
| mandible_2, replicate 2 | 3820083 | 3673342 |

**Table S3.** Genes with ASE in each tissue

| Tissue | FDR<0.05 | FDR<0.1 |
|---|---|---|
| Brain | 714 | 963 |
| Flank skin | 1103 | 1424 |
| Eyes | 918 | 1194 |
| Liver | 1139 | 1423 |
| Ventral toothplate (1) | 1606 | 2088 |
| Dorsal toothplate (1) | 1533 | 1986 |
| Mandible (1) | 1349 | 1739 |
| Ventral toothplate (2) | 1228 | 1590 |
| Dorsal toothplate (2) | 1213 | 1579 |
| Mandible (2) | 1318 | 1603 |

**Table S4**. Pairwise overlap of genes with ASE between tissues (percentages in parentheticals correspond to [number of genes with ASE in both tissues]/[total Genes with ASE in across tissues])

| | Brain | Eyes | Mandible | Liver | DTP | VTP | Flank skin |
|---|---|---|---|---|---|---|---|
| Brain | | | | | | | |
| Eyes | 376 (29%) | | | | | | |
| Mandible | 314 (18%) | 365 (19%) | | | | | |
| Liver | 270 (17%) | 340 (20%) | 368 (18%) | | | | |
| DTP | 329 (21%) | 380 (21%) | 777 (44%) | 371 (19%) | | | |
| VTP | 343 (21%) | 394 (22%) | 676 (36%) | 391 (20%) | 665 (37%) | | |
| Flank Skin | 304 (20%) | 370 (22%) | 606 (33%) | 372 (20%) | 568 (36%) | 611 (32%) | |

**Table S5**. GO terms with biased directionality in single tissue analysis

| Tissue | Biased term[1] | Permutation *P* |
|---|---|---|
| Eyes | methyltransferase activity (10/10) | 0.0018 |
| Mandible 1 | peptidase activity (22/27) | 0.0145 |
| Mandible 1 | fatty acid metabolic process (10/11) | 0.0041 |
| Flank skin | endoplasmic reticulum (38/50) | 0.0039 |

[1]In parentheses are the number of terms in the same direction out of total

**Table S6**. GO terms with biased directionality in combined test of dental tissue

| GO Term | # across tissues[1] | Permutation *P* | Fisher's combined *P* |
|---|---|---|---|
| canonical Wnt signaling pathway | 9/10 | 0.0078 | 0.0012 |
| inflammatory response | 10/12[2] | 0.0095 | 0.0086 |
| embryonic viscerocranium morphogenesis | 13/16 | 0.0093 | 0.02 |

[1]Genes in the same direction/the total number of genes with directionality

[2]In inflammatory response, one gene (*mpx*) switches direction between tissues (favoring the opposite allele)

**Table S7**. Canonical Wnt signaling genes with ASE in dental tissues

| Gene Name | Pathway | Known role in tooth development |
|---|---|---|
| *frzb* | negative regulation of canonical Wnt signaling pathway (GO:0090090); canonical Wnt signaling pathway (GO:0060070) | Yes[2] |
| *lmbr1l* | negative regulation of canonical Wnt signaling pathway (GO:0090090) | |
| *gsk3ab* | negative regulation of canonical Wnt signaling pathway (GO:0090090) | Yes[3] |
| *dab2* | negative regulation of canonical Wnt signaling pathway (GO:0090090) | |
| *sfrp2* | negative regulation of canonical Wnt signaling pathway (GO:0090090); canonical Wnt signaling pathway (GO:0060070) | Yes[4] |
| *dkk2* | negative regulation of canonical Wnt signaling pathway (GO:0090090) | Yes[5] |
| *csnk1a1* | negative regulation of canonical Wnt signaling pathway (GO:0090090) | |
| *ctnnb1* | canonical Wnt signaling pathway (GO:0060070) | Yes[6] |
| *wnt5a* | canonical Wnt signaling pathway (GO:0060070) | Yes[7] |
| *dixdc1a* | canonical Wnt signaling pathway (GO:0060070) | |
| *net1* | canonical Wnt signaling pathway (GO:0060070) | Yes[8] |
| *mllt10* | canonical Wnt signaling pathway (GO:0060070) | |
| *dvl1a* | canonical Wnt signaling pathway (GO:0060070) | |
| *dvl3a* | canonical Wnt signaling pathway (GO:0060070) | |
| *wnt2* | canonical Wnt signaling pathway (GO:0060070) | |

**Table S8**. Genes associated with term “embryonic viscerocranium morphogenesis” with ASE in dental tissues.

| Gene symbol | Gene Name |
|---|---|
| *sar1b* | secretion associated, Ras related GTPase 1B |
| *ece1* | endothelin converting enzyme 1 |
| *dlx1a* | distal-less homeobox 1a |
| *eif3ea* | eukaryotic translation initiation factor 3, subunit E, a |
| *dlx3b* | distal-less homeobox 3b |
| *gfpt1* | Glutamine--fructose-6-phosphate transaminase 1 |
| *itga5* | integrin, alpha 5 (fibronectin receptor, alpha polypeptide) |
| *nbas* | NBAS subunit of NRZ tethering complex |
| *sphk2* | sphingosine kinase 2 |
| *tmem165* | transmembrane protein 165 |
| *hmgcrb* | 3-hydroxy-3-methylglutaryl-CoA reductase b |
| *furina* | furin (paired basic amino acid cleaving enzyme) a |
| *nr3c1* | nuclear receptor subfamily 3, group C, member 1 (glucocorticoid receptor) |
| *polr1a* | RNA polymerase I subunit A |
| *ponzr1* | plac8 onzin related protein 1 |
| *gnptab* | N-acetylglucosamine-1-phosphate transferase subunits alpha and beta |

**Table S9.** Genes overlapping EcoPeaks[1]

| | ASE genes | All genes with expression | Total Genes |
|---|---|---|---|
| NE Pacific EcoPeaks (specific) | 611 | 1201 | 1625 |
| NE Pacific EcoPeaks (sensitive) | 1534 | 3382 | 4736 |

[1]Genes are considered overlapping if the reported EcoPeak interval and gene overlap by 1 or more bp; overlap with NE “specific” peaks are discussed in main text

**Table S10.** SNP density bin sizes for comparisons between ASE and CSS Z-scores

| Cuts | Number with sufficient genes[1] | Average number of genes | $P^2$ |
|---|---|---|---|
| 200 | 200 | 66 | 3.72E-10 |
| 400 | 376 | 33 | 5.38E-16 |
| 500 | 430 | 26 | 1.09E-18 |
| 1000 | 400 | 13 | 1.59E-17 |

[1]Bins with at least 5 ASE genes and background genes

[2]*P*-values are based on a Wilcoxon signed-rank test comparing bin medians (ASE vs. background); all bins were significant via permutation test ($P$<0.001).
